# upSPLAT: Early-Barcoded Library Preparation for Cost-Effective Population-Scale Genomics

**DOI:** 10.64898/2026.05.09.723775

**Authors:** Amanda Raine, Ryan J Daniels, Jonas Kjellin, Ann-Christin Wiman, Ulrika Liljedahl, Jon Ramsell, Christopher Wheat, Karl Gotthard, Mats Pettersson, Leif Andersson, Jessica Nordlund

**Author notes:** equal contribution (Ryan J Daniels, Jonas Kjellin).

## Abstract

Advances in high-throughput sequencing have substantially reduced sequencing costs, yet library preparation remains a major financial and logistical bottleneck, particularly for high-throughput applications or low-quality DNA inputs. Here, we introduce upscaled Splinted Ligation Adapter Tagging (upSPLAT), a library preparation strategy that combines early sample barcoding with single-stranded splinted ligation to enable highly multiplexed pooled sequencing at substancially reduced cost. upSPLAT supports flexible high-plex pooling and reduces per-sample library preparation costs by approximately 10-fold compared to conventional workflows. By leveraging single-strand ligation, upSPLAT is compatible with a wide range of DNA inputs, including degraded, damaged or denatured double stranded DNA, bisulfite or enzymatically converted DNA, and viral single-stranded DNA. We present two complementary workflows and evaluate their performance across multiple species and DNA qualities, demonstrating robust demultiplexing, uniform sample representation, and low barcode cross-assignment. Together, upSPLAT provides a scalable, cost-effective solution for sequencing-based studies requiring large sample numbers while preserving individual-level information.

## Introduction

Advancements in high throughput sequencing technologies have substantially reduced sequencing costs, enabling large-scale genomic studies (1). However, library preparation remains a major financial bottleneck, particularly for studies requiring high sample throughput, such as population genomics, biodiversity research, and environmental monitoring (2–6). These fields depend on sequencing large numbers of samples to characterize genetic diversity, evolutionary dynamics, and ecological adaptation (7–9), yet are often constrained by limited resources.

Traditional whole-genome sequencing (WGS) library preparation requires individual processing of each sample and is therefore costly and labor-intensive, limiting scalability. To mitigate these constraints, several alternative strategies have been developed. Reduced-representation approaches, such as restriction site–associated DNA sequencing (RAD-seq) (10) and genotyping-by-sequencing (GBS) (7,8), reduce sequencing costs by targeting only a subset of the genome. While these methods enable high sample throughput, they capture only a small fraction of genomic variation (10,11) and introduce systematic biases, including restriction fragment bias, restriction site heterozygosity, PCR-induced GC bias, and non-random haplotype sampling (12,13).

Tagmentation-based approaches using in-house Tn5 transposase have been widely adopted as more affordable alternatives to traditional workflows (14,15). However, Tn5-based protocols are sensitive to the DNA-to-Tn5 ratio, perform poorly with degraded DNA, and are prone to sequence-dependent biases.

Pooled sequencing (Pool-seq) provides a cost-effect alternative for genome-wide estimation of allele frequencies across large populations without individual-level sequencing (9). However, it sacrifices individual-level information, precluding the inference of individual genotypes, haplotypes, relatedness, and inbreeding coefficients (9,16,17). In addition, technical factors including ambiguous mapping in repetitive or structurally complex genomic regions, GC bias, base-calling errors, and PCR stochasticity, inflate variance in read counts, particularly for low-copy templates. These effects are exacerbated under conditions of low sequencing depth and in large pools, where stochastic amplification differences during early PCR cycles (e.g. early-cycle drift) are indistinguishable from true allele frequency variation (18).

Commercial methods incorporating early barcoding and pooling, such as Twist FlexPrep (Twist Bioscience), Twist 96-plex (former RipTide) (19) and AgriPrep (SeqWell) are emerging and reduce library preparation and consumable costs. Despite their convenience, these kits remain costly at scale. For studies involving large sample numbers, particular for small to midsize genomes, the library preparation costs can exceed sequencing costs.

Current library preparation strategies do not fully meet the combined demands of biodiversity and population genomics, which require scalable, low-cost processing of large sample sets while accommodating low-quality DNA. In practice, these studies often rely on low-biomass or degraded material from field collections or extreme environments (20). Here, we introduce upscaled Splinted Ligation Adapter Tagging (upSPLAT), a library preparation strategy that integrates early sample barcoding and pooling to reduce reagent and consumable costs to ∼5 USD per sample while enabling scalable processing of large sample sets. Originally developed for single-stranded DNA ligation (21–23), SPLAT is compatible with diverse DNA inputs, including degraded material. We present two pooled upSPLAT workflows and evaluate their performance across multiple species and sample types.

## Results

### Two SPLAT workflows for early barcoding and pooling

SPLAT library preparation is based on the ligation of adapters with short random overhangs (“splints”) to single-stranded DNA (ssDNA), or to double-stranded DNA (dsDNA) following denaturation. Originally developed for ssDNA, SPLAT preserves native fragment ends and is compatible with a wide range of DNA inputs, including ssDNA, dsDNA, nicked and damaged DNA. To enable affordable and scalable library preparation, we present adaptations of the SPLAT workflow that incorporate early sample barcoding and pooling using two complementary approaches that reduce overall library costs, decrease plastic and consumable usage, and increase flow-cell pooling capacity for a range of different DNA inputs.

The first approach, upSPLAT-ligation (upSPLAT-L) (Fig. 1a), introduces a 6-bp inline sample barcode to fragmented gDNA via a SPLAT adapter using ligation. This workflow operates directly on naturally fragmented or pre-fragmented DNA. Here, barcoding is applied at the 5′ end, although the workflow can be readily adapted for 3′-end or dual-end barcoding. After SPLAT ligation, which requires no end-repair, samples are pooled, and all subsequent purification, limited-cycle PCR amplification/indexing, and quality control steps are performed in bulk. upSPLAT-L is compatible with diverse DNA inputs, including degraded or low-quality samples.

**Fig. 1.**
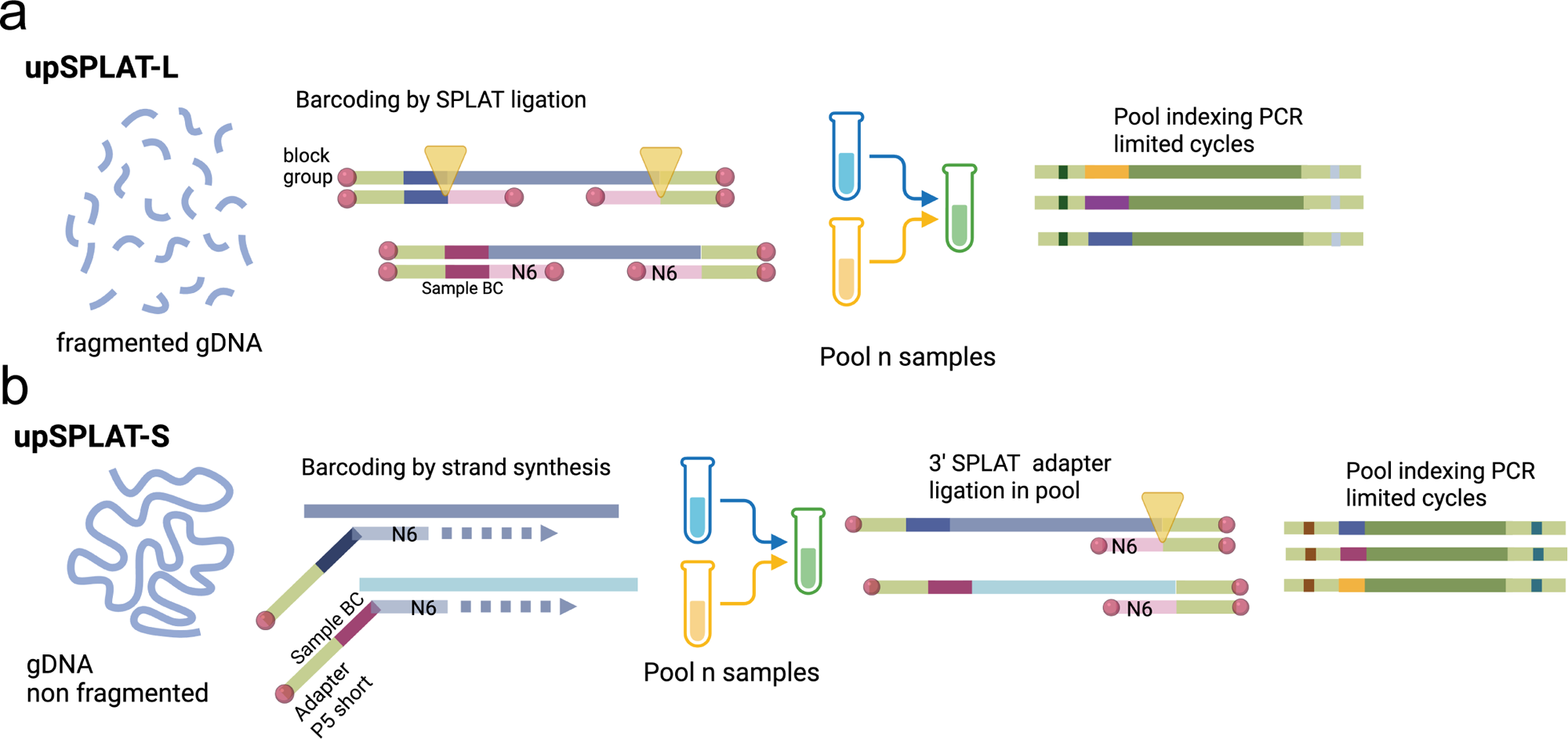
Overview of upSPLAT library preparation workflows. **(a)** In upSPLAT-L (barcoding by ligation), genomic DNA (gDNA) is first fragmented, and a barcoded 5′ SPLAT adapter together with a standard 3′ SPLAT adapter are ligated in a single reaction, preserving the true fragment ends. PCR amplification, quality control (QC), and quantification are then performed on pooled samples. **(b)** In upSPLAT-S (barcoding by strand synthesis), strand synthesis and sample barcoding are performed directly from gDNA using random hexamer oligonucleotides containing an inline barcode and a partial P5 sequence. A 3′ SPLAT adapter is subsequently ligated to pooled samples, followed by PCR amplification, QC, and quantification.

The second approach upSPLAT-strand synthesis (upSPLAT-S) (Fig. 1b), introduces a 10-bp inline barcode during strand synthesis (21,24,25). In contrast to upSPLAT-L, this workflow does not require fragmentation and is well suited for intact or high-quality DNA. Here, a random-priming oligonucleotide containing both the barcode and a partial P5 adapter-sequence is used to copy genomic DNA strands using Klenow (exo–). The low processivity and limited strand-displacement activity of this DNA polymerase generate fragments of suitable length for short-read sequencing (<1,000 bp), thereby eliminating the need for physical or enzymatic fragmentation prior to library preparation. Following strand synthesis, samples are pooled and the second adapter is attached by bulk 3′-end SPLAT ligation, after which the pooled libraries are PCR-amplified and indexed using a limited number of cycles.

By employing upSPLAT-based pooled library preparation, the cost per library can be reduced by 78 - 91% for 16-plex upSPLAT-L and -S pools, respectively compared with a conventional commercial single-sample library preparation (Table S1). Costs can be further reduced using higher multiplexing levels in the pools.

In summary, upSPLAT-L encodes sample identity during adapter ligation using barcoded adapters, preserves true fragment ends and is broadly compatible with diverse DNA inputs, whereas upSPLAT-S introduces barcodes during strand synthesis using a simple barcoded oligonucleotide, which simplifies barcode generation, facilitates high-plex pooling (as single barcode oligos are used as opposed to barcoded adapter) and eliminates the need for a separate fragmentation step.

### Demultiplexing and Pooling Performance

The pooling performance of the two protocols was evaluated by sequencing 31 pools (8–16-plex) prepared from genomic DNA of varying quality from six sample types spanning a range of genome sizes and GC content. The pools included samples from three Lepidoptera species (*Pararge aegeria*, *Lasiommata megera*, and *Parnassius apollo*), two fish species (*Clupea harengus* [herring] and *Micromesistius poutassou* [blue whiting]), *Saccharomyces cerevisiae* (baker’s yeast), and the ZymoBIOMICS Microbial DNA Community Standard (a predefined mixture of ten microbial species) (Table 1, Table S2–S3).

**Table 1.** Samples and pools included in the evaluation of upSPLAT protocols.

| Type | Short name | Species | DNA quality | N Pools upSPLAT-L | N Pools upSPLAT-S | N samples |
| --- | --- | --- | --- | --- | --- | --- |
| Herring | - | *Clupea harengus* | Mixed | 1 | 1 | 32 |
| Blue Whiting | BW | *Micromesistius poutassou* | Mixed | 6 | 10 | 243 |
| Lepidoptera | LM | *Lasiommata megera* | High | 1 | 1 | 16 |
| Lepidoptera | PA | *Parnassius apollo* | Low | 2<sup>a</sup> | - | 12 |
| Lepidoptera | PAEG | *Pararge aegeria* | High | 5 | - | 80 |
| Baker's Yeast | - | *Sacharomyces cerevisiae* | High | 1 | 1 | 1<sup>b</sup> |
| Zymo Microbial Standard | ZMB | *Listeria monocytogenes* | High | 1 | 1 | 1<sup>b</sup> |
|  |  | *Pseudomonas aeruginosa* |  |  |  |  |
|  |  | *Bacillus subtilis* |  |  |  |  |
|  |  | *Escherichia coli* |  |  |  |  |
|  |  | *Salmonella enterica* |  |  |  |  |
|  |  | *Lactobacillus fermentum* |  |  |  |  |
|  |  | *Enterococcus faecalis* |  |  |  |  |
|  |  | *Staphylococcus aureus* |  |  |  |  |
|  |  | *Saccharomyces cerevisiae* |  |  |  |  |
|  |  | *Cryptococcus neoformans* |  |  |  |  |
| Species mix | - | *Homo sapiens* | High |  |  |  |
|  |  | *Sacharomyces cerevisiae* | High |  |  |  |
|  |  | *Equus caballus* | High |  |  |  |
|  |  | *Clupea harengus* | Mixed |  |  |  |

<sup>a</sup>duplicate pools prepared for the same samples
<sup>b</sup>samples in pools are technical replicates

Across all 31 pools, a high proportion of reads were assigned to barcodes following demultiplexing (mean 98.5 ± 0.8 % and 96.7 ± 0.8 % for upSPLAT-L and upSPLAT-S respectively) (Fig. 2a, Table S4). A single outlier upSPLAT-S pool (BW-S1) showed suboptimal barcode assignment (69.5 %) and was therefore excluded in Fig. 2a but were included in the downstream analysis of pool eveness (Fig. 2b, c).

**Fig. 2.**
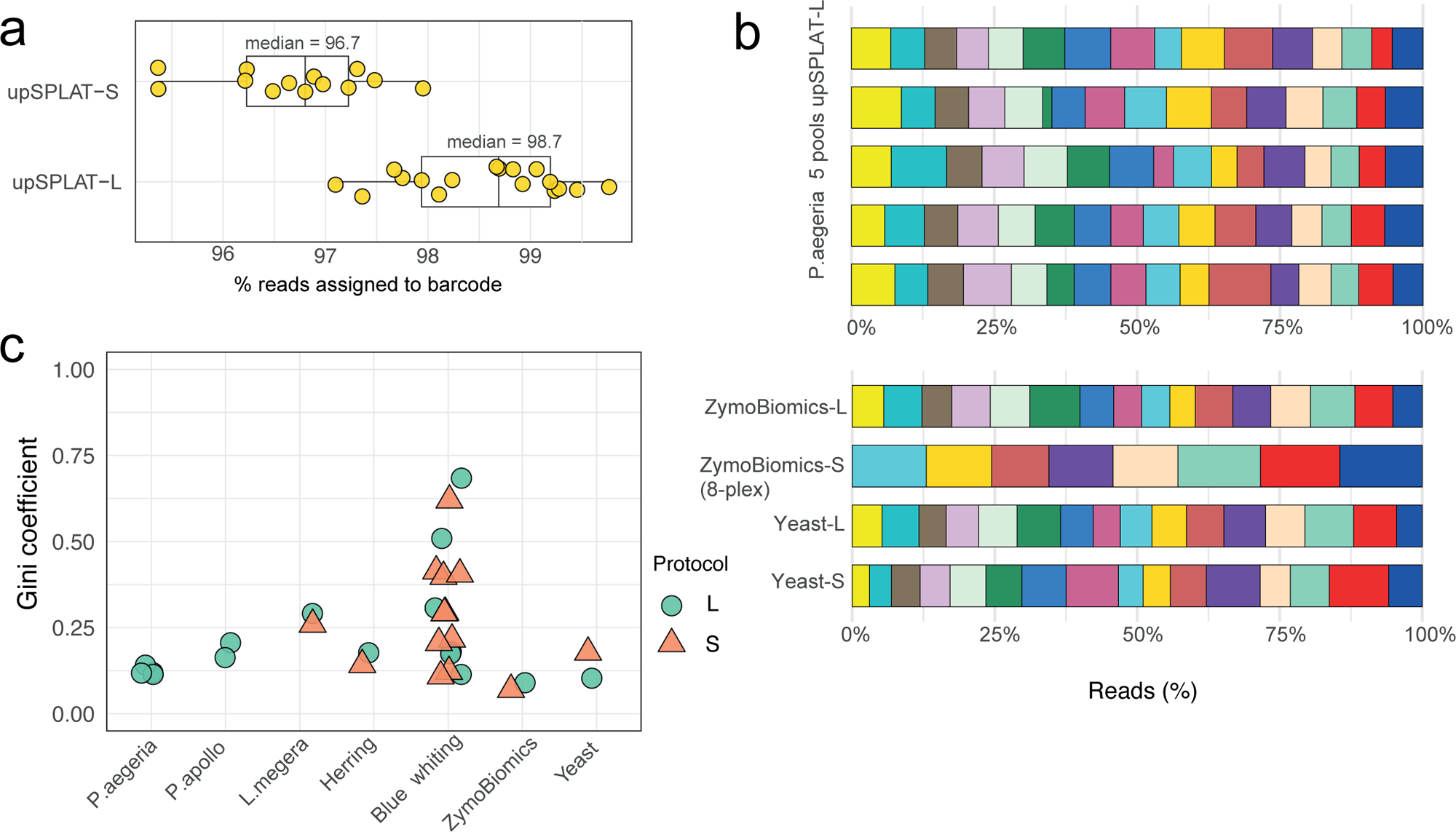
Demultiplexing performance of upSPLAT pools. (**a**) Percentage of reads assigned to expected barcodes across upSPLAT-L (n = 13) and upSPLAT-S (n = 17) pools. A single outlier upSPLAT-L pool with suboptimal barcode assignment (69.5%) was excluded from this plot. (**b**) Visualization of read distribution in a subset of demultiplexed upSPLAT pools. Stacked bar plots show the number of reads per sample within each pool. The upper panel shows five pools of 80 *Pararge aegeria* samples prepared using the upSPLAT-L protocol. The lower panels show examples of upSPLAT-L and upSPLAT-S pools consisting of technical replicates of a microbial reference standard (ZymoBIOMICS) and yeast (*Saccharomyces cerevisiae*). (**c**) Gini coefficients summarizing the distribution of reads per sample within each pool.

We quantified sample representation within the pools (pool evenness) using the Gini coefficient (G) (26), where 0 denotes a perfectly even distribution and values approaching 1 indicate increasing imbalance. Pools were generally even, with mean Gini coefficients of 0.22 ± 0.16 and 0.27 ± 0.15 for upSPLAT-L and upSPLAT-S, respectively (Fig. 2b,c; Table S5), with no significant difference between protocols (paired Wilcoxon signed-rank test, V = 10, p = 0.63). However, we observed increased unevenness in a subset of blue whiting pools. Variation in pool evenness was primarily associated with inaccuracies in DNA normalization prior to library preparation, for example due to very low sample concentrations (Fig. S1). In contrast, degraded DNA did not impair pool balance when DNA concentrations could be reliably quantified (Fig. S1). These observations indicate that pooling performance is robust across both methods, while highlighting the importance of accurate DNA quantification.

### Evaluation of barcode cross-contamination

Accurate demultiplexing of pooled libraries assumes that barcode leakage between samples during library preparation is minimal. To investigate the extent of barcode leakage within pools, we prepared mixed-species upSPLAT pools comprising genomic DNA from yeast, human, horse, and herring. Demultiplexed reads were mapped to a concatenated reference genome containing all included species (Fig. 3a).

**Fig. 3.**
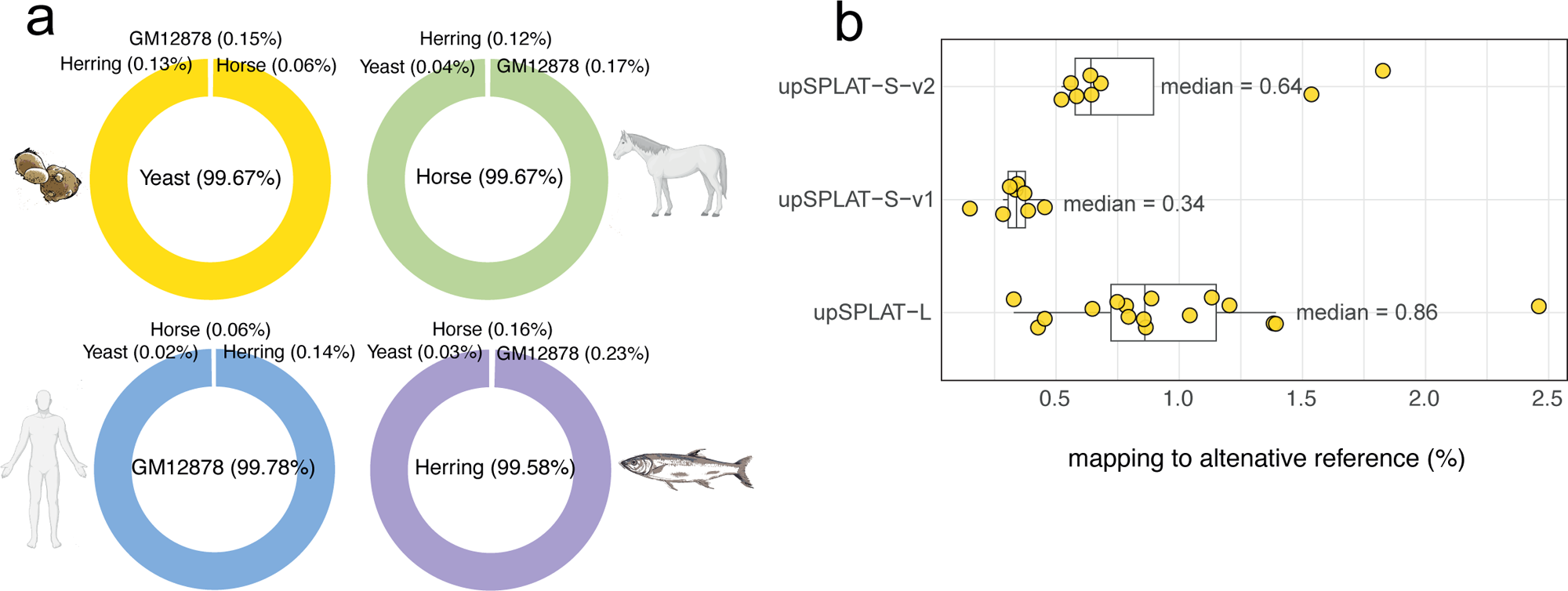
Evaluation of barcode cross-contamination. **(a)** Example of mapping specificity in samples from a mixed-species upSPLAT-S pool (v1), showing the percentage of reads mapping to the correct and alternative reference genomes for yeast, horse, human (GM12878), and herring samples. **(b)** Percentage of reads mapping to alternative reference genomes across all samples in three mixed-species upSPLAT pools. The upSPLAT-S v1 and v2 pools differ only in cleanup stringency, with v1 including two consecutive bead cleanups prior to PCR amplification, whereas v2 includes a single cleanup.

For upSPLAT-L, one mixed-species pool was generated (16-plex; four replicates per DNA sample; G = 0.131). For upSPLAT-S, two mixed-species pools (v1 and v2) were generated (8-plex; two replicates per DNA sample; G = 0.19 and 0.24 for v1 and v2, respectively). To evaluate the effect of cleanup stringency, upSPLAT-S-v1 included an additional bead cleanup step (i.e., two consecutive cleanups) prior to PCR amplification, whereas v2 included only a single cleanup at this step.

Across pools, the majority of mapped reads (median >99%) were correctly assigned to the expected species (Fig. 3b). The small fraction of misaligned reads is likely attributable to ambiguous mapping in conserved genomic regions rather than barcode leakage (Table S6). The more stringent cleanup procedure (upSPLAT-S-v1) resulted in a modest improvement in correct species assignment accuracy, reducing incorrectly mapped reads by 37% (median of 0.34 % ± 0.08 vs 0.87 % ± 0.48 in upSPLAT-S-v2; Fig. 3e). For upSPLAT-L, the mean fraction of reads mapping to an incorrect reference genome was 0.86 % ± 0.29, indicating similarly low levels of cross-assignment. These values approach those reported for index hopping on Illumina sequencing platforms (27).

Together, these results demonstrate that both upSPLAT workflows enable even pooling and acurate demultiplexing with low barcode mis-assignment, supporting reliable sample pooling following appropriate DNA quantification and normalization.

### Library performance assessed using a microbial community standard

Library quality and reproducibility were assessed using one pool each of upSPLAT-L and upSPLAT-S prepared from technical replicates of the ZympBIOMICS microbial community DNA standard with defined composition and diverse GC conent (Fig. 4a). Performance was summarized using the Measurement Integrity Quotient (MIQ) score, where higher values (maximum = 100) indicate closer agreement with the expected composition and lower bias. In the upSPLAT-L pool, observed read proportions closely matched the expected composition, including high-GC taxa, yielding a median MIQ score of 91.5 (mean = 91± 2.1; Fig. 4b). For upSPLAT-S, the corresponding value was 80 (mean 80 ± 2.6), indicating overall good performance but increased bias relative to upSPLAT-L (Fig. 4b). Notably, in upSPLAT-S, the GC-rich genome of *P. aeruginosa* was underrepresented (mean ± SD = 7.61 ± 1.30% vs expected 12%; Wilcoxon, p < 0.001; Table S7), consistent with GC-associated bias. Chimeric read levels were low (mean 1.54% ± 0.16 for upSPLAT-S and 0.72% ± 0.20 for upSPLAT-L; Table S3).

**Fig. 4.**
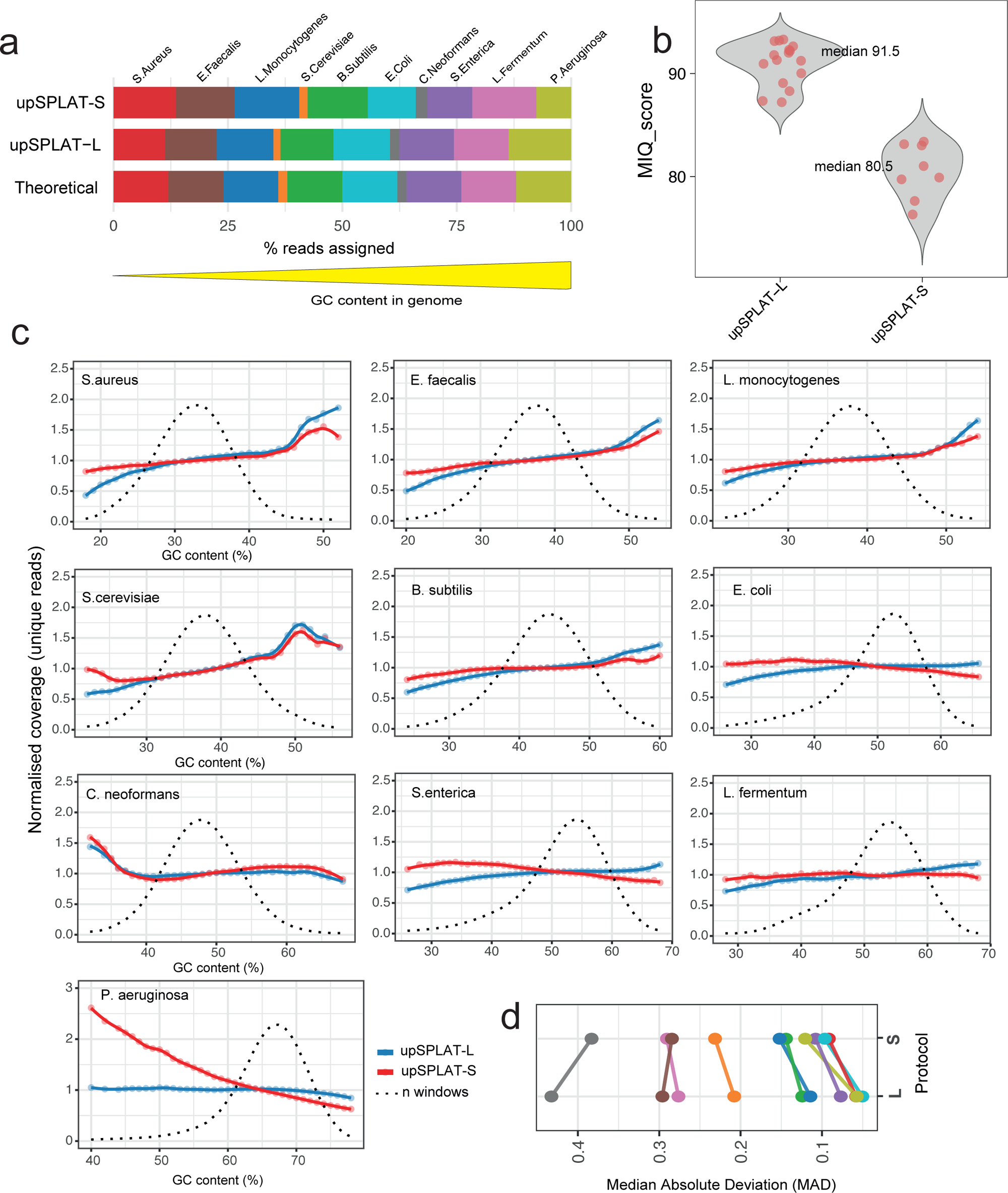
Performance evaluation of upSPLAT protocols using a microbial community standard. **(a)** Percentage of reads mapped to each species in the microbial community standard, aggregated across technical replicates within each upSPLAT pool. Both protocols show close agreement with the theoretical composition. **(b)** Distribution of Measurement Integrity Quotient (MIQ) scores for individual technical replicates across the two upSPLAT protocols, shown as violin plots. **(c)** Normalized coverage as a function of GC content for the microbial community standard, aggregated across technical replicates within each upSPLAT pool. Only GC bins representing ≥ 0.1% of genomic windows are shown. The dotted line indicates the GC content distribution of the reference genomes. **(d)** Within-sample coverage variability for the microbial community standard, measured as the median absolute deviation (MAD) across genomic windows. Genomes are color-coded as in (**a**), and MAD values are compared between upSPLAT protocols.

Coverage bias across GC content was largely similar between protocols, except for *P. aeruginosa*, where upSPLAT-S showed reduced coverage in regions below 60% GC, while upSPLAT-L maintained uniform coverage (Fig. 4c). This suggests that random-primed strand synthesis in upSPLAT-S may introduce some bias in high-GC genomes.

To complement the GC-bias analysis, overall within-sample coverage variability was quantified using the median absolute deviation (MAD) of log2 coverage-ratio values across non-overlapping genomic bins (∼5 kb), as calculated by cnvkit (28). Because this metric is influenced by genome sequence characteristics (e.g., GC content, repeats, and mappability), comparisons were performed within species to assess relative differences between protocols. Overall, the MAD values were similar between the two protocols. However, 8/10 species in upSPLAT-S libraries exhibited slightly higher coverage variability (paired Wilcoxon signed-rank test, V = 44.5, p = 0.046). (Fig. 4d, Table S8).

### Library performance assessed using upSPLAT pools from other species

Next, we analyzed GC bias and coverage variability in pools containing *L. megera*, *S. cerevisiae* (upSPLAT-L and -S), and *P. apollo* (upSPLAT-L). Consistent with the microbial sample analysis, GC bias was slightly more pronounced and variable in upSPLAT-S compared to upSPLAT-L (Fig. S3).

For *L. megera* replicates, MAD values were very similar across protocols (mean 0.248 ± 0.008 and 0.274 ± 0.016 for upSPLAT-L and upSPLAT-S, respectively; Table S8). In contrast, for *S. cerevisiae* technical replicates, higher coverage variability was observed in upSPLAT-S libraries (mean 0.361 ± 0.026 vs 0.083 ± 0.002 for upSPLAT-S and upSPLAT-L, respectively; Table S8) (paired Wilcoxon signed-rank test, V=136, p= 0.0002)

Library complexity was further assessed using duplicate read rates and Preseq complexity (c_curve) analysis. Non-optical duplicate levels were consistently low (mean 4.21 ± 2.38%), whereas optical duplicate rates varied substantially across libraries (10–40%; mean 23.9 ± 11.9%) and increased with mean fragment length within pools, possibly reflecting suboptimal loading concentrations for pools with longer inserts on the NovaSeq X (Table S3). Preseq complexity curves indicated high library complexity for most samples (Fig. S3).

Overall, both upSPLAT protocols produced libraries of good quality and reproducibility across technical replicates and diverse genome compositions. upSPLAT-L provided more uniform coverage across genomes and samples, supporting robustness for complex and compositionally diverse inputs, whereas upSPLAT-S performed satisfactorily for most samples although exhibited moderately higher variability in coverage uniformity across species and samples.

### Comparison of upSPLAT protocols with Tn5 libraries

Next, we compared upSPLAT with Tn5-based library preparation (17). Genomic DNA from 32 Baltic herring (*Clupea harengus membras* L., 1758) samples was sequenced with the two upSPLAT protocols (upSPLAT-L, individuals =1-16; upSPLAT-S, individuals = 17-32). These samples had previously been subjected to low-pass whole-genome sequencing (∼1x) using a Tn5-based protocol.

The 16-plex pools were sequenced to mean coverages of 9x ± 3 (upSPLAT-L) and 17x ± 4 (upSPLAT-S). Pool evenness within both pools was comparable to previous pools (G = 0.14 and G = 0.17 for upSPLAT-L and upSPLAT-S, respectively; Fig. 5a). To enable direct comparison with the Tn5 data, reads from the demultiplexed upSPLAT libraries were downsampled following alignment and deduplication. After downsampling, all three protocols exhibited similar coverage distributions and comparable mean coverage values (Table S9), differing by less than 3% from the Tn5 data.

**Fig. 5.**
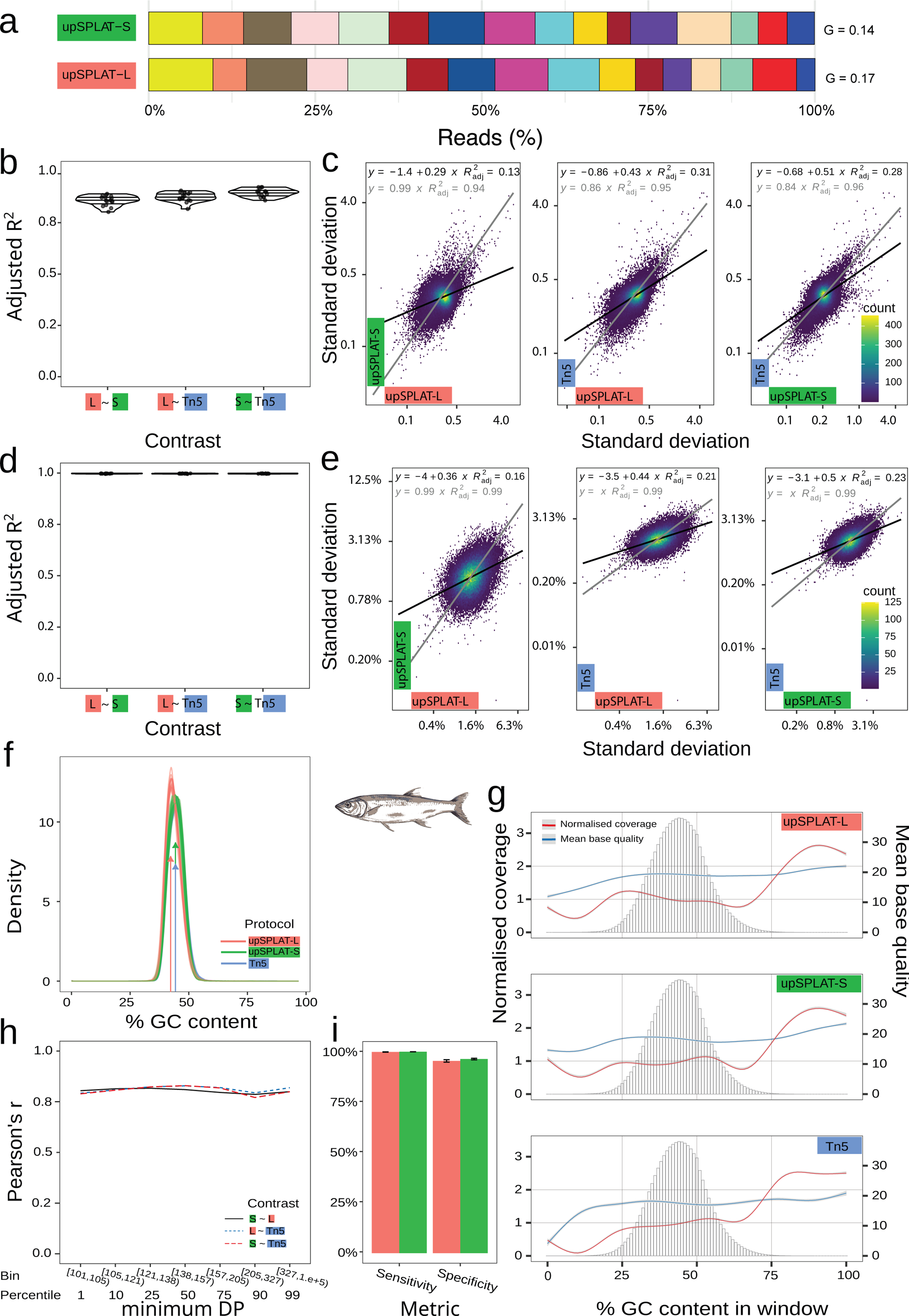
Coverage uniformity, GC bias, and variant concordance across upSPLAT-S and upSPLAT-L protocols. **(a)** Read distribution in herring upSPLAT-S and L pools. **(b, d)** Pairwise correlations of coverage and GC content for the downsampled data. In **(b)** distribution of adjusted R^2^ estimates for pairwise correlations of mean coverage in 10 Kb windows and in **(d)** GC content among protocols. **(c, e)** Pairwise correlations of standard deviation of coverage **(c)** and GC content **(e)** estimated per 10 Kb window from all 16 samples. Best fit linear regression line indicated for free y-intercept (black) and fixed y-intercept (grey). Adjusted R^2^ (proportion of variance explained) are shown in the scatter plot. Note that the axes are on log2 scale. To aid visualisation, data have been binned on both the x- and y-axes. Density presented here as the number of data points within the respective bin. **(f)** Percent GC estimated by 10 Kb window for each library. Arrows indicate the approx. peak of the distributions. **(g)** Changes in coverage and base quality across the % GC content distribution for downsampled data. Loess regression curves fit for normalised coverage (red) and mean base quality score (blue). Error margin indicated for each line in faded buffer. Histogram shows the frequency distribution of reads across GC content bins. Values estimated for 100 bp windows. **(h)** Changes in Pearson’s correlation coefficient of allele frequencies between protocols with increasing coverage. Correlations were performed using only data from the respective minimum DP (depth of coverage) bin **(i)** Specificity and sensitivity of the upSPLAT protocols compared to data generated with a standard Illumina method. Results are based on overlapping SNPs between the respective data sets. The data are from pooled high coverage resequenced samples from (29).

Pairwise correlations of mean coverage calculated over 10-kb windows were high across protocols (adjusted R^2^ > 0.9, p < 0.001 for most comparisons; Fig. 5b). The strongest concordance was observed between upSPLAT-S and Tn5 libraries (adjusted R^2^ = 0.86–0.92), a pattern that was also evident when non-downsampled data were compared (Fig. S4).

Additionally, coverage variability was assessed by examining concordance in the standard deviation of windowed coverage across samples, in order to determine whether the same genomic regions exhibited elevated variability across protocols. Correlations were high between protocols (adjusted R^2^ = 0.94–0.96; p < 2 x 10⁻¹⁶), indicating largely consistent patterns of region-specific variability (Fig. 5c). Concordance was highest for upSPLAT-S versus Tn5 comparisons and slightly lower for upSPLAT-S versus upSPLAT-L comparisons.

All three library preparation protocols produced similar genome-wide GC content distributions (Fig. 5d). Pairwise correlations of GC content across protocols were consistently high (adjusted R^2^ > 0.99, p < 0.001; Fig. 5e), indicating near-identical GC representation. A minor skew toward lower GC content was observed for upSPLAT-L libraries (Fig. 5f), resulting in a slightly lower median GC estimate compared to the Tn5 and upSPLAT-S libraries (46.0% versus 47.2%). Variability in GC content, assessed as the standard deviation across genomic windows, was also highly correlated among protocols (adjusted R^2^ = 0.99, p < 2 × 10⁻¹⁶; Fig. 5e), suggesting that GC-related variation was primarily driven by sequence context rather than protocol-specific effects.

Mean normalized coverage profiles as a function of GC content were broadly similar across all protocols (Fig. 5g). Coverage declined in low-GC regions, most strongly for Tn5 libraries, while both upSPLAT protocols retained higher coverage in these regions. All protocols showed increased coverage at GC-rich extremes. Base quality metrics were largely comparable across protocols for GC contents between 25% and 75% (where the majority of windows are) (Fig. 5g).

We next assessed genotype concordance across upSPLAT and Tn5 protocols using an allele frequency–based framework. Allele frequencies were estimated from the 16 individuals sequenced with each upSPLAT protocol (non downsampled data) and from 150 randomly selected individuals from the Tn5 dataset to achieve comparable effective coverage. Allele frequency estimates were highly concordant across all protocol comparisons and sequencing depth (DP) bins (Fig. 5h and Fig. S5). Correlations strengthened with increasing depth, rising from an adjusted R^2^ of 0.79 in the lowest depth bin ([101,105), 1st percentile) to 0.82 in intermediate DP bins ([157,205), 75th percentile). At the highest depth bins, concordance declined slightly (adjusted R^2^ ≈ 0.79), likely reflecting enrichment of repetitive or structurally complex regions (e.g., tandem duplications), where mapping ambiguity increases and protocol-specific differences become more pronounced.

Furthermore, SNP detection performance was evaluated by comparing variants identified using the upSPLAT protocols to those detected from a pooled whole-genome resequencing dataset (30x coverage, 50 individuals) from the same population (Hästskär, Sweden) prepared with a different library preparation (standard Illumina kit) (29). This resequenced data serves as a reference set of true SNPs (here after ‘*defacto SNPs’*) that are polymorphic in the population (polymorphic *defacto SNPs*). The total number of polymorphic SNPs detected differed between protocols, with 172,073 SNPs identified by upSPLAT-L and 518,394 by upSPLAT-S, reflecting differences in sequencing depth (average 9x vs 17x, for upSPLAT-L and upSPLAT-S respectively). Of the total polymorphic SNPs detected by the upSPLAT protcols, 68% (116,566 loci) for upSPLAT-L and 81% (422,291 loci) for upSPLAT-S overlapped with SNPs detected in the pooled resequencing dataset. Within this shared set, agreement between protocols was high. Both upSPLAT workflows showed similarly high sensitivity (99.8-99.9%; false-negative rate = 0.002–0.001) and high specificity (95.4-96.3%; false-positive rate = 0.045–0.037; Fig. 5i), indicating robust and consistent SNP detection performance.

Despite lower coverage and fewer individuals, the proportion of polymorphic SNPs relative to the total number of defacto SNPs detected by upSPLAT (41.1% and 38.9%, for L and S respectively) was only slightly lower than in this resequenced dataset (50.2% of 16,158,082 SNPs detected at 30× coverage).

Together, these results suggest that both upSPLAT protocols maintain coverage profiles comparable to conventional Tn5-based library preparation for herring DNA, and that protocol-specific biases are minor. Moreover, allele frequency estimates and SNP detection obtained using upSPLAT-L and upSPLAT-S are highly concordant with each other and with alternative library prep methods, supporting the suitability of upSPLAT for accurate genotype-level analyses.

## Discussion

Early sample multiplexing and pooling during library preparation enables substantial reductions in reagent and consumable costs. Reduced plastic usage is also important from a sustainability perspective, and as sequencing scales continue to grow, minimizing environmental impact becomes increasingly relevant.

Here, we present upSPLAT, a flexible library preparation strategy that integrates early barcoding with pooled processing. Across the conditions tested, both upSPLAT workflows supported robust demultiplexing, low barcode cross-assignment, and generally even sample representation, while substantially reducing per-sample costs.

The two workflows differ in their performance characteristics and potential applications. upSPLAT-L provides more uniform coverage across GC content and is compatible with a wide range of DNA inputs, including degraded and single-stranded DNA, supporting its use in challenging sample types. However, for high molecular weight DNA it requires fragmentation prior to library preparation, which increases per-sample cost. In contrast, upSPLAT-S is more streamlined and cost-efficient. Although it showed increased sequence-dependent bias, particularly for high-GC genomes, likely introduced during random-primed strand synthesis (30,31) it is well suited for large sample series where balancing sequencing and library preparation costs is critical. Primer design for strand synthesis can be further tailored to GC- or AT-rich genomes (19) to improve performance across diverse genomic contexts.

Both upSPLAT workflows can be scaled beyond the pooling levels evaluated here. In Table S10, we provide sequences for 64 barcode oligonucleotides for upSPLAT-S, enabling higher degrees of multiplexing. However, based on our results, moderate multiplexing (up to 16-plex) provides a practical balance between cost efficiency, robustness, and experimental flexibility. At this level, pooling reduces the impact of sample-to-sample variability while minimizing the risk that poorly normalized samples dominate sequencing output. Using this strategy, 96 samples can be processed in six pools, substantially reducing consumables and labor while enabling high throughput (>1500 samples per sequencing lane when combined with standard (96 UDI) indexing strategies.

In summary, upSPLAT provides a cost-effective and scalable alternative to conventional library preparation, enabling efficient processing of large sample sets while preserving individual-level resolution. The method is particularly well suited for applications requiring flexible handling of diverse DNA inputs and moderate-to-high sample throughput. The workflows are amenable to automation and are well suited for large-scale population genomics, biodiversity and environmental genomics, agrigenomics, and museomics.

## Methods

### Samples

ZymoBIOMICS Microbial DNA standards were obtained from Zymo Research. Yeast (*Sacharomyces Cerevisiae*) gDNA was purchased from Sigma Aldrich. Horse gDNA was purified from immune cells using the Qiagen AllPrep Kit. Baltic herring (*Clupea harengus* subsp. *membras* L. 1758) were sampled in 2017 from Hästskär on the Swedish east coast (60.5345 N, 17.7323 E). Tail fin tissue was taken from freshly caught fish and stored in 98% ethanol at – 80 °C until needed. DNA was extracted with the DNeasy 96 Blood & Tissue Kit (Qiagen). All DNA samples were quantified using Quant-iT or Qubit.

### Library preparation

Detailed step-by-step upSPLAT protocols are (at submission) available on protcolos.io. The general outline of the library preparation procedures are outlined here.

#### upSPLAT-S

Normalized gDNA (50 ng in 5 μl) in 96-well plates were prepared. Strand synthesis oligos (SSOs) were diluted to 50 nM in 96-well plates. The general sequence of the SSOs were 5’AmMC6/TTCCCTACACGACGCTCTTCCGATCTX10N6 (IDT), where X10 = 10 bp sample index (Table S10 and S11) and N6 = random hexamer. SSOs (2ul) were added to samples and the DNA/primer mixes were heated at 95 ℃ for 3 minutes and then placed on ice for 5 minutes. A strand synthesis master mix (comprising Klenow exo-, 1x Blue Buffer and dNTPs) was dispensed to each well (8 μl), the plate vortexed, spun down briefly and incubated at 37 ℃ for 45 minutes. The strand synthesis reactions were then quenched by adding 25 µl of AMPure XP beads to each well. Reactions (n=16) were pooled into a 2 ml Eppendorf tube. After washing beads two times with 80 % ethanol the pooled DNA was eluted in in 20 µl H_2_O. T4 DNA ligation buffer (0.5 µl of 10X buffer) and Thermolabile Exo I (0.5 μl) (New England Biolabs) was added and incubated 37 ℃ for 4 minutes. A second bead purification was performed by adding 30 µl H_2_O and 50 µl AMPure beads and eluted in 16 µl H_2_O. The pool was the heat denatured for 3 minutes at 95 ℃, followed by immediately placing the tubes on ice for 5 minutes. Ligation of 3’ SPLAT adapters (Table S10) were then performed in a total volume of 40 µl, by adding 6 µl of 3’ SPLAT adapters (100 µM stock), 4 µl of 10x T4 DNA Ligase buffer, 10 ul of PEG8000 (40%) followed by thorough vortexing for 1 min before adding 1 µl PNK and 2 µl T4 DNA Ligase HC (30 U/µl, ThermoFisher). The ligation reaction was vortexed thoroughly and incubated 20 minutes at 20 ℃ followed by 20 minutes at 37 ℃. The ligation reaction was then purified by adding 60 µl of water and 90 µl of AMPure beads and finally eluted in 15 µl H_2_O. Amplification of the pools for 4-5 cycles were carried out with KAPA HiFi PCR mix and NEB UDI index primers (10 µl) in a final volume of 50 µl. The final library pools were purified with a 0.7X bead ratio and library profiles checked on Tape Station. If adapter dimers were still visible a second bead purification was performed.

#### upSPLAT-L

DNA samples were normalized by concentration (to 10 ng/ul) and fragmented to 400-550 bp using a Covaris E220. For degraded DNA samples fragmentation was omitted. 50 ng per sample in 9 µl was dispensed into 96-plate wells together with 1 µl of ET-SSB (2x diluted) and the plate was incubated in a thermocycler at 95 ℃ for 3 minutes and then immediately placed on ice for 5 minutes. Barcoded 5’ SPLAT adapters (n=16) and the standard non-barcoded 3’-end SPLAT adapter were diluted to 40 uM (from a 100 uM stock plate) and 1 µl of each adapter added to wells columnwise. SPLAT adapter oligos sequences are found in Table S10. A ligation master mix was prepared (per sample: 2 µl 10x Ligation Buffer, 5 µl 40% PEG8000, 1 µl PNK (10 U/µl), 0.25 µl ATP (100 mM), 1 µl T4 DNA Ligase (30 U/ µl), 0.75 µl H_2_O) and thoroughly mixed by vortexing before adding 10 ul to each well. Plates were vortexed, spun down and placed in a thermocycler at 20 ℃ for 20 minutes followed by 37 ℃ for 20 minutes. Ligation reactions were quenched by adding 25 AMPure beads/well and were then pooled, AMpure bead purified twice and finaly eluted in 15 ul H_2_O. The pools were amplified for 4-5 cycles using KAPA HiFi PCR mix and NEB UDI index primers. The final libraries were purified with a 0.7 X bead ratio and library profiles checked on Tape Station. If adapter dimers were still visible a second bead purification was performed.

### Pool demultiplexing and read alignment

Demultiplexing of pooled reads was performed with the PAMLD algorithm in Pheniqs (v2.1.0) (32). Reads were assigned to samples based on inline barcodes of 10 bp (upSPLAT-S) or 6 bp (upSPLAT-L) using a 0.95 confidence threshold. Pool evenness was assessed using the Gini coefficient (*G*), calculated with the Gini function from the *ineq* R package. Values range from 0 to 1, where lower values indicate a more even distribution of reads across samples.

Read alignment and initial quality control were performed with nf-core/sarek pipeline v 3.5.1 (33). The pipeline was run with the following parameters: --skip_tools baserecalibrator --trim_fastq true --trim_nextseq 10. It should be noted that Klenow-based strand synthesis may generate a terminal 3′ A overhang, therefore optional 3′ trimming may be considered for the upSPLAT-S protocol.

For the ZymoBIOMICS Microbial DNA Standards samples, reads were mapped to a concatenated reference genome containing all species in the mock community. These samples were further analyzed with the Shotgun Sequencing MIQ (Measurement Integrity Quotient) Calculator (https://github.com/Zymo-Research/miqScoreShotgunPublic) to obtain standardized quality scores (MIQ scores) for mapping and computing performance, as well as the estimated proportion of chimera-like reads.

To quantify barcode swapping, reads from single-organism samples within cross-species pools were mapped against a multi-species reference (human, yeast, herring and horse). The number of reads mapping to each distinct reference was determined using samtools view (34) filtering for primary, non-duplicate alignments with a minimum mapping quality of 10 (-c -F 3332 -q 10). All reference genome versions are detailed in Table S2.

### Library complexity and coverage uniformity analyses

Library complexity was estimated using preseq c_curve (v3.2) (35). Coverage uniformity was assessed through two metrics: GC bias calculated with Picard CollectGCBiasMetrics (v3.1.1) using 150bp window size and the Median Absolute Deviation (MAD) of coverage calculated with cnvkit (v0.9.11) (28).

To obtain species-specific metrics for the ZymoBIOMICS Microbial DNA standards samples, aligned reads from the initial sarek run (see above) were first pooled by protocol (upSPLAT-L and upSPLAT-S) and subsequently extracted per species. The species-specific reads where then aligned to respective individual genome reference using the nf-core/sarek pipeline and used as input for the coverage uniformity analyses.

### Comparison of Baltic Herring upSPLAT and Tn5 libraries

#### Alignment and Downsampling

Herring upSPLAT-L and upSPLAT-S 16-plex pools were demultiplexed as described above and reanalysed using the same procedure as for the Tn5 data. Raw reads were quality-assessed using FastQC v.0.11.9 and summarized with MultiQC v.1.9 (36). Adapter sequences were trimmed using TrimGalore! v.0.6.6 with default settings, including removal of low-quality bases (–quality 20) and exclusion of reads shorter than 20 bases (–length 20). Reads were aligned to the chromosome-level assembly for Atlantic herring (*Clupea harengus*; Ch_v2.0.2) (37) using using BWA v0.7.18 component BWA-MEM. Paired-end fastq files were split into six files of equal size using SeqKit split2 function and each subset was aligned independently. Resulting alignments were concatenated, sorted and post-processed using Samtools v1.20. Duplicate reads were marked and removed using Picard v3.1.1. Downsampling of aligned, deduplicated libraries was performed using the Samtools view -s option, with the downsampling probability *p* function defined as:

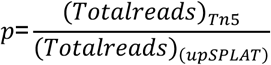

#### Performance metrics: Coverage and GC bias

For low-pass Tn5 and downsampled upSPLAT data, protocol-specific differences in genome-wide coverage were investigated using metrics from non-overlapping 10 kb windows. The distribution of mean coverage values was examined to assess variability within protocols.

Congruence between protocols was evaluated using Pearson’s correlation to assess whether windows deviated from the mean coverage in a similar direction across protocols. Samples were randomly paired across protocols to estimate correlations. Coverage estimates were obtained from aligned BAM files using MosDepth v0.3.6 (38) with the option --by 10000 to compute mean coverage per window. Post-processing was performed in R v4.3.1 using Tidyverse packages. To prevent zero inflation, particularly in the Tn5 data, windows with zero coverage were excluded prior to correlation analysis. Correlations were computed using the R package *ggpubr* and the function stat_cor(). Linear regression models were fit with the y-intercept fixed at zero based on theoretical expectations.

In addition, the correlation of coverage standard deviation across windows was assessed. For each window, the standard deviation of coverage across all samples was calculated, and adjusted R^2^ values based on Pearson’s r were computed between each pair of protocols. The y-intercept was fixed at (0,0) for regression fitting based on the theoretical expectation. In all cases, the correlation coefficient value improved substantially compared to using *c* ≠ 0.

In a manner analogous to the coverage analysis, GC content was calculated for non-overlapping 10 kb genomic windows. Congruence between protocols was evaluated using Pearson’s correlation adjusted R^2^ values based on Pearson’s r of GC content values across corresponding genomic windows. GC content was estimated using a custom script based on Bedtools v2.31.1 (39). Because GC content is an intrinsic property of the genome, this analysis was used to confirm that the genomic regions did not show protocol-specific coverage patterns. In addition, the relationship between normalized coverage and GC content was examined using the CollectGcBiasMetrics command from the Picard toolkit with default settings. The overall GC bias profile for each protocol was modeled using a generalized additive model (GAM) implemented with the gam function from the mgcv R package using the formula y ∼ s(x, bs = “cs”).

#### Allele Frequency Concordance and SNP Detection Performance

Joint genotype calling was performed using the Sentieon pipeline. The GATK HaplotypeCaller (29) was run using the following parameters: -algo Haplotyper -genotype_model multinomial -emit_mode gvcf - emit_conf 30 -call_conf 30. Genotyping calling was performed using flags: GVCFtyper using - emit_mode CONFIDENT.

Filtering was conducted using GATK VariantFiltration and SelectVariants with the following criteria:: MQ <40.0 || MQRankSum < -12.5 || ReadPosRankSum < -8.0 || QD <2.0 || FS >60.0 || DP <3 || GQ <20. Following filtering, samples from each protocol were pooled *in silico* for allele frequency estimation. Allele frequencies at each locus were calculated using a custom Python script.

To ensure reliable comparisons, additional filtering steps were applied. Loci with a total depth (DP) below 100 were removed, and loci with missing data in any protocol were excluded to retain only positions shared across datasets. Furthermore, loci with allele frequencies outside the detectable range given the sample size were excluded. For 16 diploid individuals, we conservatively consider the minimum detectable non-zero allele frequency as 2/32 alleles = 0.0625, and the maximum is 30/32 alleles = 0.9375. Allele frequencies outside this range cannot be reliably distinguished from fixation given the limited number of chromosomes sampled and therefore do not provide informative variation for correlation-based comparisons.

Congruence between protocols was assessed using Pearson’s correlation coefficient, implemented in R with the ggpubr package using the function stat_cor(). SNP calling performance was further evaluated by comparing variants identified in this study to previously published resequencing data from the same population (Hästkär) (29). Here we are interested in accurately identifying a loci as a polymorphic SNP. We use pooled resequence data with 30x coverage based on 50 Hästskär individuals. Variant calling was performed using the full dataset of 53 populations of Atlantic herring as per (29), and this represents *a priori* known SNPs for the species (*defacto* SNPs). Filtering was applied based on the genome-wide parameter distribution using the following criteria: QD <5.0 || FS >40.0 || MQ <50.0 || MQRankSum < 4.0 || ReadPosRankSum < 3.0 || ReadPosRank-Sum > 3.0 || DP <100 || DP >5400. SNPs from the pooled sequences were treated as the set of SNPs truly polymorphic within Hästskär. From this reference, true positive and false positive SNP calling rates were estimated for each protocol.

Sensitivity (true positive rate) was calculated as:

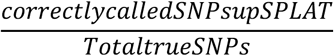

Specificity (true negative rate) was calculated as:

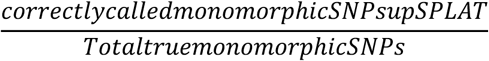

Analyses were restricted to the Hästskär population to avoid inflating false-negative estimates that would arise from comparisons against all known species-wide SNPs.

For each dataset, SNPs and indels were separated using bcftools norm -m -any | bcftools norm –m +both. Loci with no data were removed, and only genomic positions present in both datasets were retained using: bcftools isec -n=2 -c snps -w1,2 <file1> <file2>.

The upSPLAT datasets were filtered for minor allele frequency (MAF) > 0.06, while the pool dataset was filtered for MAF > 0.02 to account for differences in sample size and detection limits. Agreement between protocols was assessed using bcftools isec -c some -n +1 <file1> <file2>, which identifies loci present in either or both datasets and requires that at least one alternate allele is shared between datasets. In this framework, true negatives represent loci known to be polymorphic at the species level but correctly identified as monomorphic in the Hästskär population.

## Acknowledgments

This work was supported by a SciLifeLabTechnology Development grant. We thank the SciLifeLab National Genomics Infrastructure, SNP&SEQ Technology Platform, which is funded by the Swedish Research Council and the Knut and Alice Wallenberg Foundation, for assistance with data generation. The data handling was enabled by resources provided by the Swedish National Infrastructure for Computing (SNIC) and the National Academic Infrastructure for Supercomputing in Sweden (NAISS). SNIC and NAISS are partially funded by the Swedish Research Council through grant agreement no. 2022-06725. We thank Arianna Cocco for assistance with extraction of herring DNA samples and Anders Lundmark for assistance with computational analysis.

## Data availability

Raw sequencing reads will be available from NCBI SRA at submission.

## Supplementary Figure Legends

**Fig. S1.**
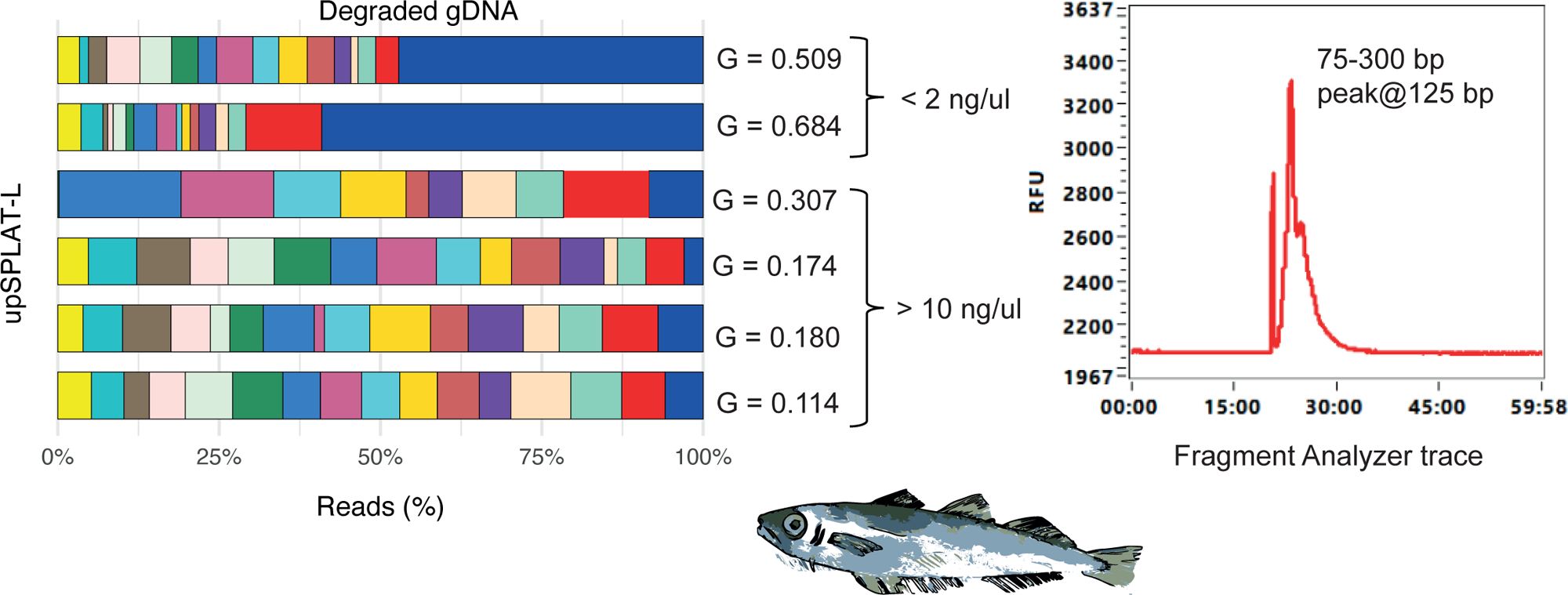
Effects of gDNA quality and input normalization on pooling evenness in blue whiting samples. Degraded blue whiting DNA samples prepared with upSPLAT-L yielded relatively even pools when input quantities allowed reliable normalization (i.e. DNA concentration >10 ng/µl), whereas very low-input samples (<2 ng/µl) resulted in skewed pools (Gini coefficient > 0.3). A Fragment Analyzer trace shows a representative example of a degraded blue whiting sample prepared using the upSPLAT-L protocol.

**Fig. S2.**
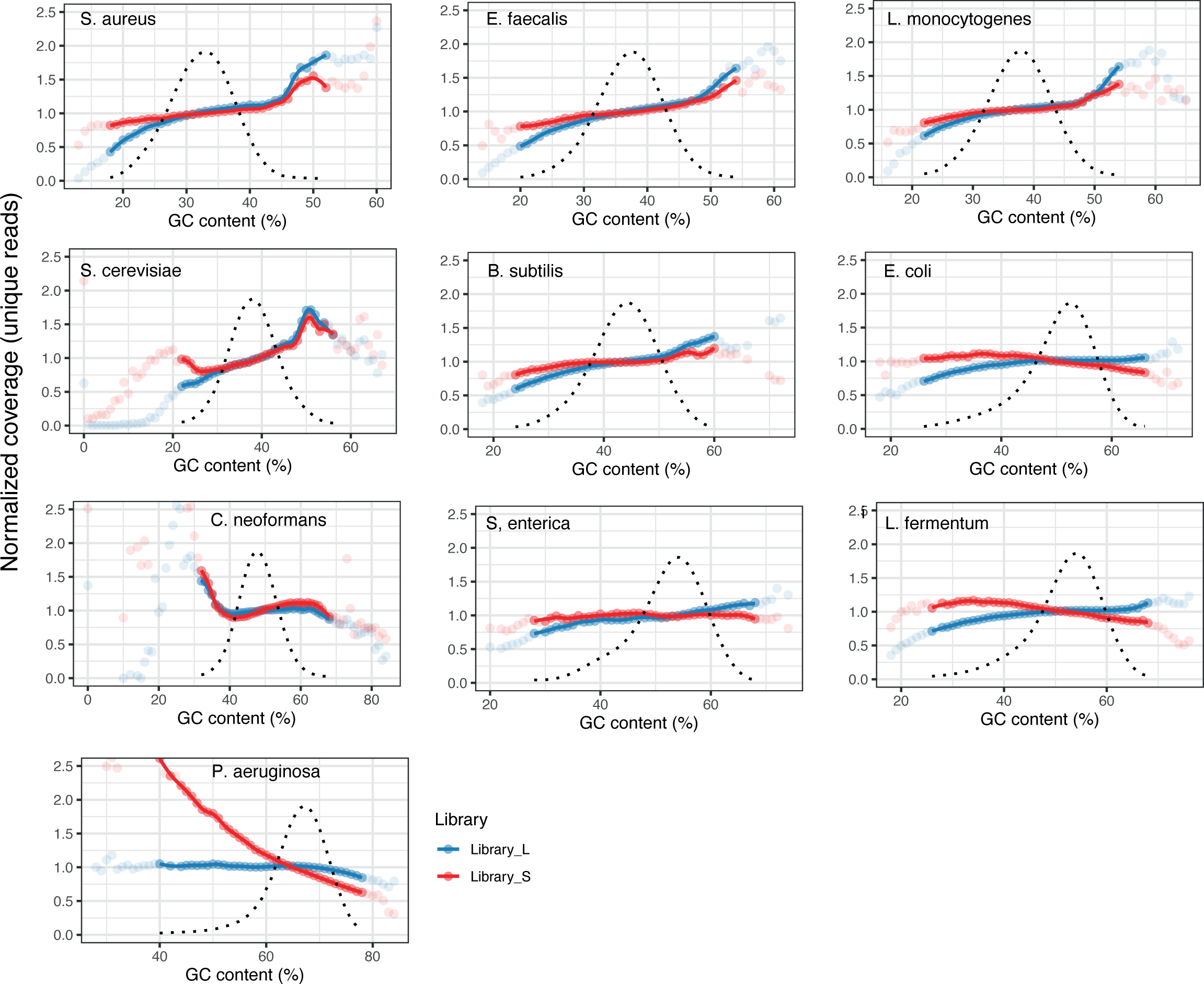
Extended GC bias plots for microbial community standard samples. Same as Fig. 2, but also visualizing bins representing <0.1% of the total windows (faded). Bins with extremely few windows (i.e., <100) are not shown.

**Fig. S3.**
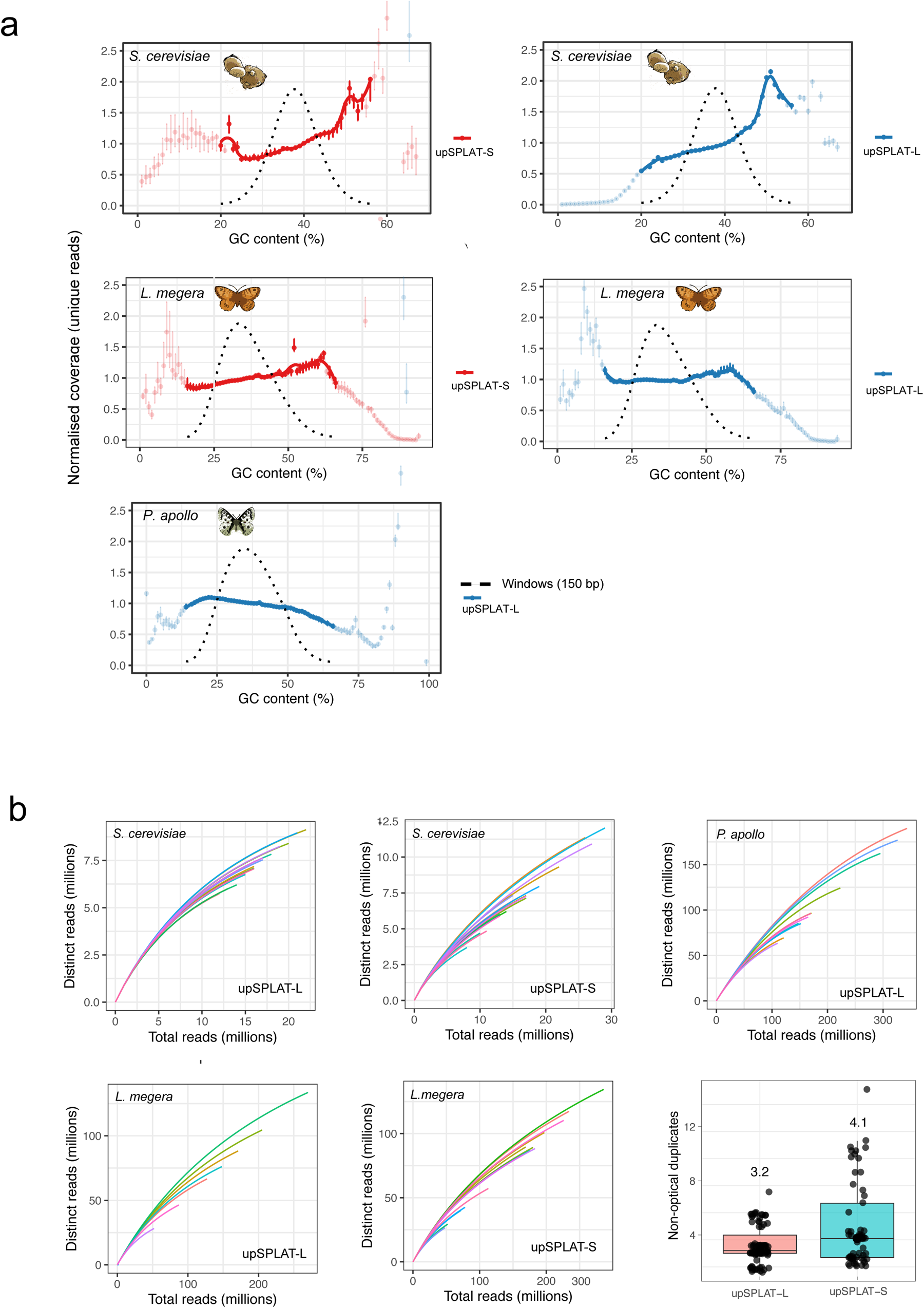
GC bias and library complexity across upSPLAT pools. **(a)** Mean normalized coverage across GC content bins (± SD) is shown for each pool. Yeast (*S. cerevisiae*) pools comprise technical replicates (n = 16), whereas *P. apollo* (n = 12) and *L. megera* pools (n = 8 for upSPLAT-L and n = 16 for upSPLAT-S) comprise biological replicates. GC bins representing <0.1% of total genomic windows are shown in faded color. Bins with very few windows (<100) were excluded. **(b)** Library complexity curves generated using the *preseq* c_curve function for the same pools and samples as in **(a).** Boxplots show the percentage of non-optical duplicates for libraries prepared with the two upSPLAT protocols.

**Fig. S4.**
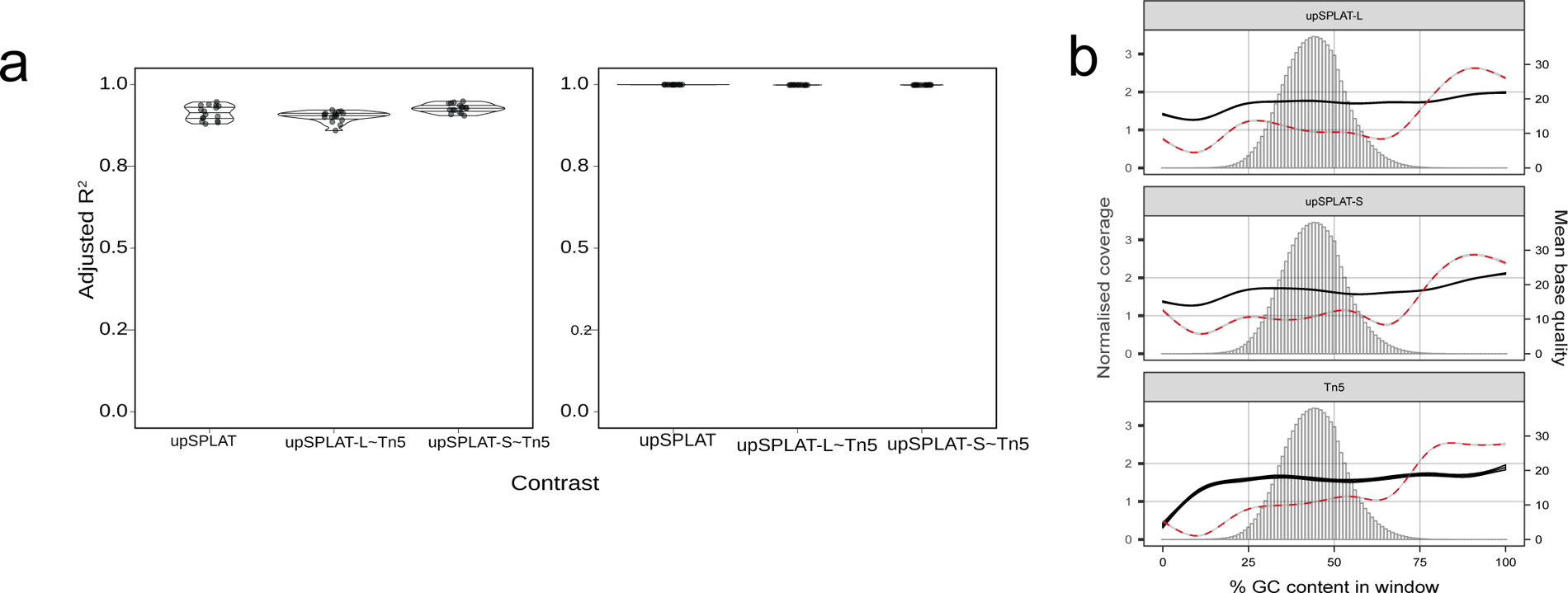
Comparison of GC content and coverage profiles across upSPLAT and Tn5 herring libraries (non-downsampled data). **(a)** Pairwise correlations of coverage and GC content for non-downsampled herring data. Distributions of adjusted R² values for pairwise correlations of mean coverage in 10 kb windows and GC content in 10 kb windows across protocols. **(b)** Coverage and base quality as a function of GC content for non-downsampled data. Loess regression curves are shown for normalized coverage (red) and mean base quality score (black), with shaded regions indicating the error margin. The histogram shows the distribution of reads across GC content bins. Values were estimated using 100 bp windows.

**Fig. S5.**
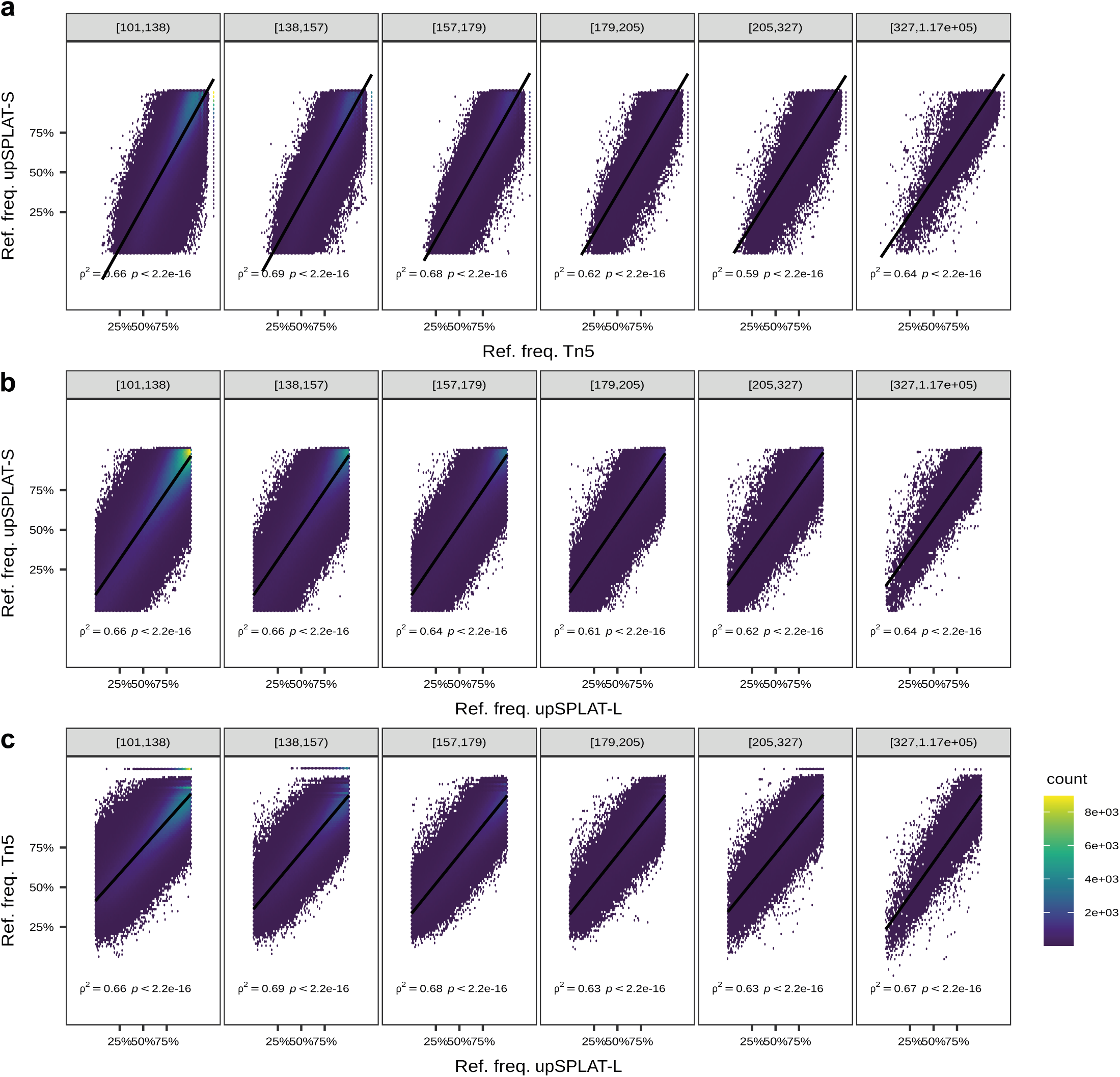
Correlation of reference allele frequency estimates between protocols made with raw upSPLAT and 1x Tn5 data. Contrasts of **(a)** upSPLAT-S against Tn5 **(b)** upSPLAT-S against upSPLAT-L **(c)** Tn5 against upSPLAT-L. Facets (left to right) show depth (DP) bins. Correlations were performed using only data from the respective DP bin. Indicated in each plot is the adjusted R^2^ and p-value as well as the best fit linear regression. Data density represented as the number of nearest neighbours to each data point (count). 150 individuals from the Tn5 protocol were pooled to achieve the equivalent coverage as the upSPLAT protocols. Note axes have arcsine transformation.

## Supplementary Tables

Table S1. upSPLAT cost estimates

Table S2. List of reference genomes

Table S3. General sample metrics

Table S4. Reads assigned to barcodes

Table S5. Gini coefficients

Table S6. Species-mix read mapping

Table S7. Percent reads assigned to species in microbial community standard pool replicates.

Table S8. MAD values obtained from cnvkit

Table S9 Herring coverage and downsampling metrics per chromosome.

Table S10. Barcode oligos

Table S11. Per sample barcode information

